# Mammalian retinal specializations for high acuity vision evolve in response to both foraging strategies and morphological constraints

**DOI:** 10.1101/2024.08.25.609608

**Authors:** Emily E. K. Kopania, Nathan L. Clark

**Affiliations:** Department of Biological Sciences, University of Pittsburgh, Pittsburgh, PA, USA

**Keywords:** vision, orbit convergence, area centralis, horizontal streak, phylogenetic comparative methods, mammals

## Abstract

Vision is a complex sensory system that requires coordination among cellular and morphological traits, and it remains unclear how functional relationships among traits interact with ecological selective pressures to shape the evolution of vision. Many species have specialized high visual acuity regions in the retina defined by patterns of ganglion cell density, which may evolve in response to ecological conditions. For example, ganglion cell density can increase radially towards the center of the retina to form an area centralis, which is thought to improve acuity towards the center of the visual field in predators. Another example is the horizontal streak, where ganglion cells are dense across the center of the retina, which is thought to be beneficial in horizon-dominated habitats. At the morphological level, many have proposed that predation selects for high orbit convergence angles, or forward-facing eyes. We tested these hypotheses in a phylogenetic framework across eutherian mammals and only found support for the association between the horizontal streak and horizon-dominated habitats. We also tested if retinal specializations evolve in response to orbit convergence angles. We found that horizontal streaks were associated with side-facing eyes, which may both facilitate panoramic vision. Previous studies observed that some species with side-facing eyes have an area centralis shifted towards the temporal side of the retina, such that the high acuity region would project forward, but this relationship had not been tested quantitatively. We found that the temporal distance of the area centralis from the center of the retina was inversely correlated with orbit convergence, as predicted. Our work shows a strong relationship between orbit convergence and retinal specializations. We find support that both visual ecology and functional interactions among traits play important roles in the evolution of some ocular traits across mammals.

## Introduction

Many species rely on visual cues for important behaviors such as detecting predators, finding food, and finding mates (Caves et al., 2018). Vision-related traits exhibit remarkable divergence across taxa, and these differences may be adaptations to different visual environments (Caves et al., 2017; Caves et al., 2018; Baker & Venditti, 2019; Nilsson, 2021). Some vision-related traits are associated with ecological conditions even after accounting for shared evolutionary history, suggesting that these traits may evolve in response to selective pressures imposed by the visual environment (Caves et al., 2017; Potier et al., 2017; Baker & Venditti, 2019; Cantlay et al., 2023; Potier et al., 2023; Caves et al., 2024; Chong et al., 2024). For example, visual acuity is associated with habitat complexity in both fish and birds (Caves et al., 2017; Caves et al., 2024), and relative corneal size increases in nocturnal mammals (Baker & Venditti, 2019). However, developmental and functional constraints are also expected to play an important role in the evolution of ocular traits (Nilsson, 2021). For many ocular traits, it remains unclear if they are associated with ecological conditions that may impose selective pressures on vision. Additionally, vision is a complex sensory system that integrates traits from the cellular level, such as the arrangements of retinal cell types, to the morphological level, such as the size and orientation of the eyes. These cellular and morphological traits may be evolving in response to the same selective pressures and could also impose functional constraints on each other, so understanding their relationships is important for gaining a complete understanding of the evolution of vision (Hughes, 1977).

One such trait is orbit convergence, or the angle of the orbits relative to the anterior-posterior axis of the skull (Figure 1A). A higher orbit convergence angle corresponds to more forward-facing eyes, with an angle of 90° indicating that the orbits face directly forwards. Orbit convergence is highly correlated with binocular overlap, or the extent to which the visual fields of each eye overlap, which is thought to facilitate visual acuity and stereopsis, or depth perception (Heesy, 2004; Read, 2021). Lower orbit convergence an-gles correspond to more sideways-facing eyes, which may facilitate panoramic vision and allow animals to have a wider field of view (Walls, 1942; Hughes, 1977; Heesy, 2007). Within mammals, primates have exceptionally high orbit convergence angles, which has been hypothesized to be an adaptation to arboreal lifestyles or nocturnal predation in the ancestral primate species (Cartmill, 1972, 1974; Heesy, 2007). Outside of primates, carnivores tend to have high orbit convergence, while groups with more herbivorous species such as lagomorphs and artiodactyls tend to have low orbit convergence (Walls, 1942; Hughes, 1977). These observations led to the hypothesis that high orbit convergence is adaptive in predators, whereas low orbit convergence is beneficial in species that experience high predation rates to facilitate predator detection across a wider visual field (Walls, 1942; Hughes, 1977). This is often assumed to be the case (Casares-Hidalgo et al., 2019), but this pattern has not been tested broadly across mammals in a phylogenetic framework. Some studies have focused on a few species or particular clades (Noble et al., 2000; Vega-Zuniga et al., 2013; Pilatti & Astúa, 2016; Smith et al., 2018; Casares-Hidalgo et al., 2019), but these groups do not encapsulate all the variation in orbit convergence that exists across mammals. Others have found that orbit convergence is associated with predation and nocturnality (Heesy, 2007) or with body mass specifically in forested environments (Changizi & Shimojo, 2008), but these studies did not correct for phylogenetic relatedness, and most of these associations are no longer significant after controlling for phylogeny (Griffin, 2017). Thus, it remains unclear if arboreality, nocturnality, predation, or some combination of these traits are associated with high orbit convergence across mammals.

**Figure 1.**
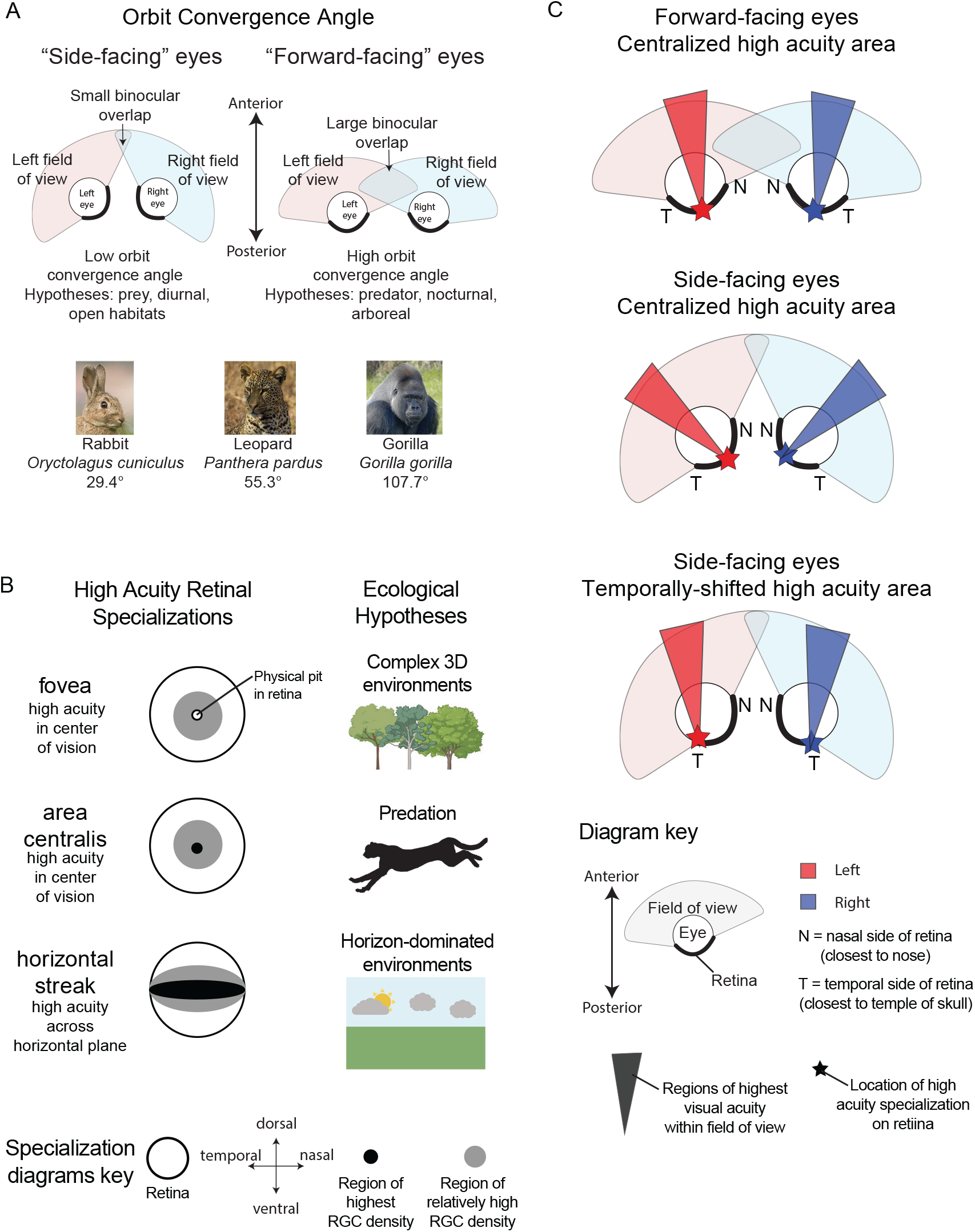
Ocular trait evolutionary hypotheses. (A) Diagrams depicting different orbit convergence angles, hypotheses for ecological conditions that may act on orbit convergence, and example species depicting the range of orbit convergence angles in mammals. Eye diagrams are shown from the dorsal perspective such that the top is the anterior direction and the bottom is the posterior direction. Red represents the left eye’s field of view, blue represents the right eye’s field of view, and the purple area represents the overlap between the two, or the binocular overlap. Rabbit photo by JJ Harrison and used under Creative Commons Attribution-Share Alike 3.0 Unported license. Leopard photo by Thomas Fuhrmann and gorilla photo by Thurundir, both used under Creative Commons Attribution-Share Alike 4.0 International license. (B) Retinal specialization diagrams and hypotheses for ecological conditions they may be associated with. Specialization diagrams are adapted from (Moore et al., 2017). RGC = retinal ganglion cell. Arboreal and horizon-dominated environments created using Biorender. Cheetah silhouette is a public domain image from PhyloPic. (C) Predicted directions of high acuity vision based on orbit convergence angle and position of high acuity retinal specialization. Species with forward-facing eyes and centrally located high acuity specializations will have the highest visual acuity towards the front (top), whereas species with side-facing eyes and centrally located high acuity specializations will have the highest visual acuity towards the sides (middle). Thus, species with side-facing eyes are predicted to have high acuity specializations located more towards the temporal side of the retina to facilitate higher visual acuity towards the front (bottom) (Collin, 1999).

Cellular phenotypes in the eye, such as specialized regions of high acuity, have also diverged across taxa and may also be associated with ecological conditions. Retinal regions of high acuity impart the ability to see fine spatial details, and they arise from localized concentrations of ganglion cells, which send the visual signal to the brain (Collin, 1999; Moore et al., 2017; Caves et al., 2018). Most vertebrates do not have consistent ganglion cell density throughout the retina, but instead have regions of high and low density; the human retina, for example, ranges over 100-fold from 200 cells/mm^2^ near the periphery to 38,000 cells/mm^2^ towards the center (Curcio & Allen, 1990). Many vertebrate taxa including mammals have evolved different high acuity retinal specializations, which are defined by patterns of high retinal ganglion cell density (Collin, 1999; Moore et al., 2017). One specialization, the fovea, is an indentation in the retina surrounded by a region of high cell density (Figure 1B). Other retinal specializations include the area centralis, a centralized region of high cell density, and horizontal streak, an elongated region of high cell density stretching across the retina (Figure 1B). These specializations have evolved repeatedly across vertebrates, suggesting they may have adaptive potential and may evolve in response to different aspects of visual ecology (Collin, 1999; Moore et al., 2017). Within mammals, only primates have a fovea (Bringmann et al., 2018), but the area centralis and horizontal streak may have evolved multiple times independently. Many have proposed hypotheses regarding the ecological drivers that select for these retinal specializations. For example, the area centralis is thought to help detect prey movement and therefore may be an adaptation for predation (Moore et al., 2017; Yoshimatsu et al., 2020). The horizontal streak is hypothesized to be associated with “horizon-dominated” environments, particularly ground-foraging species in relatively unobstructed environments such as grasslands or deserts (Hughes, 1977). Some mammal species support these predictions (Hughes, 1977; Moore et al., 2017; Navarro-Sempere et al., 2018), but these hypotheses have not been directly tested across mammals in a phylogenetic framework.

Ecological traits such as predation, nocturnality, or horizon-dominated environments may impose selective pressures on both orbit convergence and high acuity retinal specializations. For example, a horizontal streak in combination with low orbit convergence may facilitate panoramic vision (Hughes, 1977). High orbit convergence and the fovea are both thought to improve stereopsis, or depth perception, which may be beneficial for predators to detect prey (Cartmill, 1972; Steenstrup & Munk, 1980; Heesy, 2004; Moore et al., 2017; Bringmann, 2019). These traits may also impose functional or developmental constraints on each other. Species with low orbit convergence (i.e., side-facing eyes) and an area centralis may be more likely to have the area centralis located towards the temporal side of the retina, closer to the temple of the skull. This temporal shift of the area centralis in species with side-facing eyes is predicted to facilitate high acuity vision towards the front, despite the side-facing orbits [Figure 1C; (Hughes, 1977; Collin, 1999; Moore et al., 2012)]. While some studies have found support for these predictions in particular species (Collin, 1999; Moore et al., 2012), it remains unclear if orbit convergence is correlated with the types and position of retinal specializations across mammals.

Here, we test if orbit convergence and retinal specializations are associated with ecological traits, and with each other, through a phylogenetic comparative study across eutherian mammals. We used the most complete mammal phylogeny to date based on genomic data (Genereux et al., 2020), measured orbit convergence from museum specimens, and combined these data with published data on orbit convergence, retinal specializations (Supplemental Table S1), and ecological traits (Wilman et al., 2014) to address four main questions:

1. Which ecological traits are associated with orbit convergence after controlling for phylogeny
2. Is the area centralis associated with predator spe-cies?
3. Is the horizontal streak associated with horizon-domi-nated environments?
4. Is orbit convergence higher for species with a centralized retinal specialization (fovea or area centralis) and lower for species with a horizontal streak or temporally shifted area centralis?

## Materials and Methods

### Samples and data collection

We measured orbit convergence from museum specimens from the Carnegie Museum of Natural History for 28 mam-mal species (Supplemental Table S1) using the method from (Ross, 1995; Noble et al., 2000). Briefly, we used a custom-built dihedral goniometer to measure the angle between the plane of the orbit and the sagittal plane (Supplemental Figure S1). We stabilized skulls using clay or rice and measured the vertical distance from the base of the goniometer to the inion, basion, and prostion of the skull to ensure the sagittal plane was parallel with the bottom of the goniometer. We then folded the top part of the goniometer and adjusted the position of the skull until three pins of equal length touched three points on the orbit. For some specimens, we used the orbitale superius, orbitale anterius, and orbitale inferius to mark the plane of the orbit (Ross, 1995; Heesy, 2004). For other specimens, there was no orbitale inferius or it was difficult to identify, so we used the orbitale posterius instead (Casares-Hidalgo et al., 2019) (Supplemental Table S1). We then used a digital protractor (Insize CO., LTD) to measure the angle between the two planes of the goniometer. To collect data from species representing a wide range of sizes, we had three sizes of goniometers built (3 inches^2^, 12 inches^2^, and 36 inches^2^). We also obtained orbit convergence measurements from another 63 species from published sources (Heesy, 2004, 2005; Casares-Hidalgo et al., 2019) for a total of 91 species with orbit convergence data.

We used information on retinal specializations from published studies (Supplemental Table S1). Some studies define retinal specializations using photoreceptor density (Collin, 1999; Schiviz et al., 2008; Moore et al., 2012), but we only included studies that categorized retinal specializations based on retinal ganglion cell density for consistency, and because retinal ganglion cell density is thought to be the best predictor of visual acuity (Hughes, 1977; Collin, 1999). For species that had a fovea or area centralis, we measured the relative distance and angle of the specialization from the center point of the retina by implementing the method from (Moore et al., 2012) in Fiji (Schindelin et al., 2012)(Supplemental Figure S2). We manually marked the outermost points of the retinal topographic maps on the dorsal, ventral, nasal, and temporal axes, and marked the center of the retinal specialization. We then used built-in ImageJ macros to calculate the minimum bounding circle of the retina, the center point of the retina, the radius of the retina, and the distance and angle of the retinal specialization from the center point. We normalized the specialization distance by the retina radius, and then used trigonometry functions to calculate the temporal distance of the retina from the center using the specialization distance and angle.

We used ecological and body mass data from the EltonTraits 1.0 dataset (Wilman et al., 2014). Species were considered “predators” if invertebrates and vertebrates were greater than or equal to 70% of their diets, excluding scavengers. Species were considered “herbivores” if greater than or equal to 70% of their diets were plant material. For activity pattern, we categorized species as “light” (exclusively diurnal), “dark” (exclusively nocturnal or nocturnal and crepuscular), or “mix” (any other combination of activity patterns). For foraging strategy, we added a “fossorial” category and changed *Heterocephalus glaber, Fukomys damarensis, Ellobius lutescens*, and *Ellobius talpinus* from “ground” to “fossorial” since they are known to spend the majority of their lives underground and do not have fully developed eyes (Partha et al., 2017). We considered species to be in “unobstructed, horizon-dominated” habitats if they primarily reside in grassland, savannah, or tundra biomes (Myers et al., 2024) and excluded small rodents that are unlikely to experience unobstructed fields of view even in these open habitat types (Supplemental Table S1). We used the Zoonomia mammal phylogeny (Genereux et al., 2020). In some cases, we used orbit convergence and retinal specialization data from a closely-related species when data were not available for the species in the Zoonomia dataset (Supplemental Table S1). In these cases, we used the ecological data for the same species as the ocular trait data. We also used ecological data from congeneric species if none were available from the same species as the ocular trait data (Supplemental Table S1).

### Phylogenetic analyses

We visualized traits on the Zoonomia phylogeny using *phytools* V2.1-1 (Revell, 2012). We also used *phytools* to calculate phylogenetic signal using the function *phylosig* with 1000 simulations. To test for an association between orbit convergence and ecological conditions, we used phylogenetic generalized least squares (PGLS, (Martins & Hansen, 1997)) implemented in the R package *nlme* V3.1-164 (Pinheiro J, 2020) with a Pagel’s λ approach to model the underlying phylogenetic correlation structure. We used the log_10_ of body mass to control for potential allometric scaling relationships (Changizi & Shimojo, 2008; Cantlay et al., 2023; Potier et al., 2023) and ran our models both with and without body mass as a covariate. Because the activity pattern trait had multiple categories, we used a post-hoc Tukey test to compare means among categories using the *glht* function in the package *multcomp* V1.4-25. To test for differences in orbit convergence among species with different types of retinal specializations, we used the same PGLS approach. We also used a simulation-based phylogenetic ANOVA (Garland et al., 1993) using the *phytools* function *phylANOVA* with 10,000 simulations (Revell, 2012).

To test the relationships between retinal specializations and ecological conditions we compared the likelihood ratios between dependent and independent models of discrete trait evolution using a maximum likelihood estimation approach in BayesTraits V4.0.1 (Pagel & Meade, 2006). We encoded each trait as a binary variable to test each of our hypotheses (e.g., horizontal streak present or absent, ground forager or not) and corrected for multiple tests across these different hypothesis tests using Benjamini-Hochberg correction. All statistical tests and R packages were implemented using R version 4.3.2.

### Data availability

Orbit convergence data that we measured from museum specimens are available in Supplemental Table S1. Sources from the literature for orbit convergence and retinal specializations are in Supplemental Table S1 and in the supplemental material under Supplemental References. Scripts for ImageJ macros and phylogenetic analyses are available on GitHub (https://github.com/ekopania/mammal_retinas).

## Results

### No support for associations between ecological conditions and orbit convergence

We first evaluated the evolution of orbit convergence across 91 placental mammals through a combination of published data (Heesy, 2007; Casares-Hidalgo et al., 2019) and new measurements from museum specimens (Figure 2A). To verify the accuracy of our method, we measured orbit convergence for nine species that also had values reported in previous studies, and we found that our measurements were consistent with those reported in the literature (Supplemental Figure S3). We found that phylogenetic signal was relatively high, with Pagel’s λ = 0.78, where λ=0 indicates no phylogenetic signal and λ=1 represents a Brownian motion model of evolution (P < 0.001 based on a likelihood ratio test comparing λ = 0.78 to λ=0). In other words, more closely related species tended to have more similar orbit convergence values. This is consistent with observations in carnivores (Casares-Hidalgo et al., 2019), suggesting that evolutionary history may play an important role in shaping the observed values of orbit convergence among extant species both within and across major mammalian clades. Across mammals, most species with large orbit convergence angles, or more “forward-facing” eyes, are in the carnivore and primate orders (Figure 2A).

**Figure 2.**
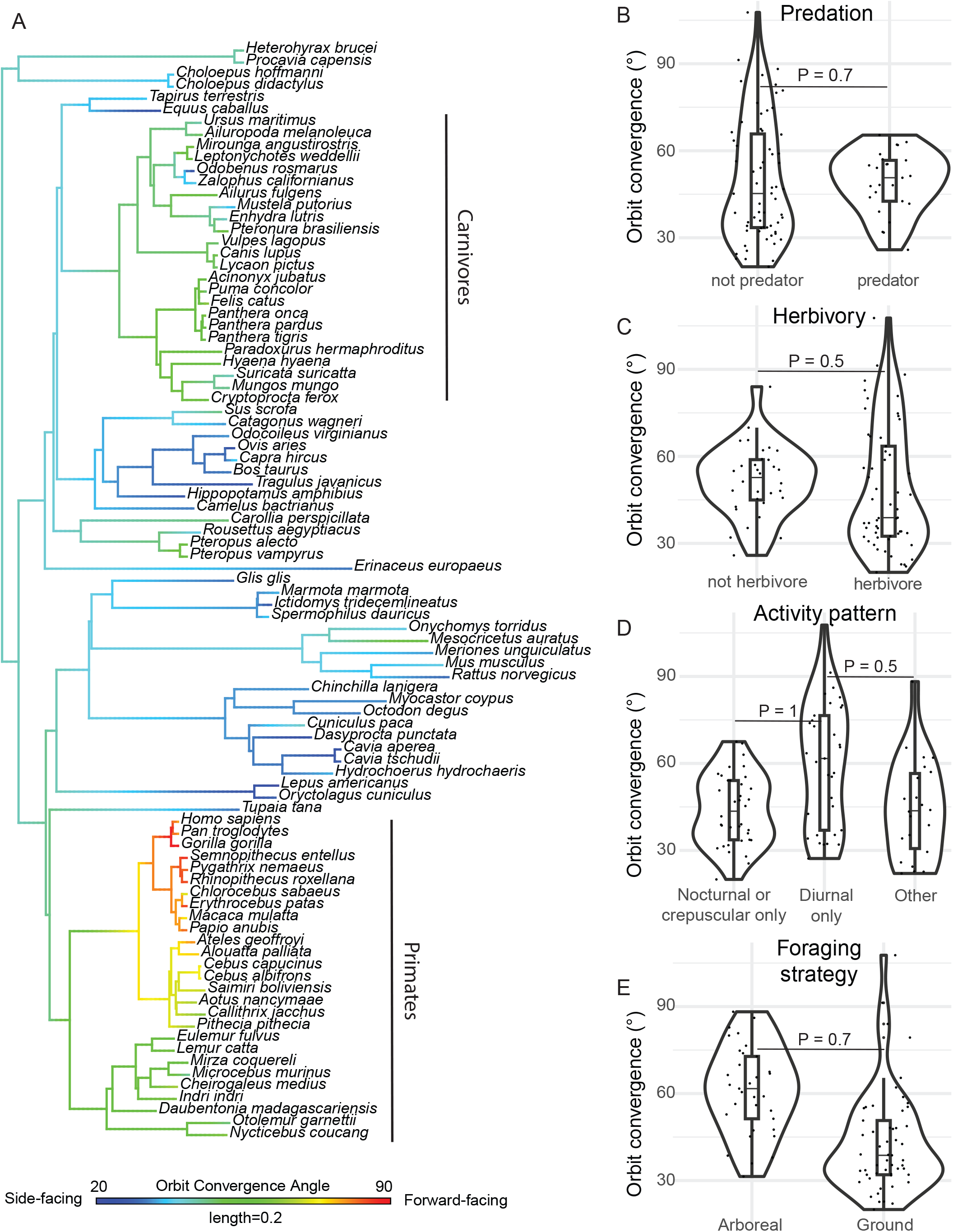
Orbit convergence evolution across mammals. (A) Orbit convergence angles overlayed on the mammalian phylogeny, with warmer colors representing higher angles or more “forward-facing” eyes. (B-E) Violin plots comparing orbit convergence angles for species in different ecological categories. Each point represents a species, and thick horizontal lines within the violin plots represent the median orbit convergence angle for species in the given ecological category. P-values are based on phylogenetic generalized least squares with body mass as a covariate.

We then asked if orbit convergence angle is associated with ecological conditions after controlling for phylogeny. Surprisingly, we did not find a significant difference in orbit convergence between predators and non-predators based on a phylogenetic generalized least squares test (P = 0.69; Figure 2B). This result held whether we included body mass as a covariate or not (Supplemental Table S2). It is difficult to quantify predation risk in a consistent way across all mammals, but species that are often considered “prey species” with low orbit convergence angles tend to be small, ground-foraging herbivores such as artiodactyls, rodents, and lagomorphs (Walls, 1942; Hughes, 1977; Heesy, 2007). We did not find statistical support for the hypothesized negative relationship between orbit convergence and herbivory (species with >70% of their diet as plant material; PGLS P = 0.48; Figure 2C). There was also no significant difference when considering an interaction between herbivory and ground foraging (PGLS P = 0.35) or herbivory and body mass (PGLS P = 0.95). Although we observed a trend towards higher orbit convergence in predators and lower orbit convergence in herbivores (Figure 2), we did not find statistical support that orbit convergence is associated with predators after phylogenetic correction.

Some have hypothesized that high orbit convergence facilitates vision in low light by improving visual acuity in the region of binocular overlap, and thus nocturnal species are predicted to have higher orbit convergence (Cartmill, 1972; Heesy, 2007; Read, 2021). However, we also did not find support for this hypothesis (PGLS P = 0.997; Figure 2D). The nocturnal predation hypothesis predicts that the combination of nocturnality and predation selects for higher orbit convergence, but there was also no significant interaction between predation and nocturnality (PGLS P = 0.75). Because high orbit convergence and binocular vision are thought to improve depth perception, orbit convergence may be higher in arboreal species to facilitate navigating in a complex, three-dimensional environment ((Changizi & Shimojo, 2008); reviewed in (Cartmill, 1974)), while lower orbit convergence may facilitate panoramic vision in ground-foraging species, which tend to experience more open or horizon-dominated environments (Hughes, 1977; Heesy, 2007). Although there was a trend towards higher orbit convergence in arboreal species compared to ground-foraging species, this relationship was not significant after phylogenetic correction (PGLS P = 0.69; Figure 2E). Overall, shared evolutionary history plays an important role in the patterns of orbit convergence across mammals, and we could not separate these effects from potential associations with ecological conditions. These results were similar whether we included body mass as a covariate or not (Supplemental Table S2).

### Ecological conditions may contribute to retinal specialization evolution

Retinal specializations are thought to be important visual adaptations to different ecological conditions, but these traits have largely been studied with a focus on individual species or a few closely related species. To understand broader patterns of retinal specialization evolution, we gathered data from the literature on retinal specializations in 82 species and overlayed these data on the mammal phylogeny (Figure 3). We found evidence that both the area centralis and horizontal streak have evolved repeatedly in mammals. Many species have both an area centralis and horizontal streak, but there are also multiple independent clades with only an area centralis or only a horizontal streak (Figure 3). Other specializations also occur in mammals, including the fovea in primates and the presence of a vertical streak or “anakatabatic area” which is common in artiodactyls (Supplemental Table S1). We focused on the area centralis and horizontal streak, because they appear to be the most common specializations and show evidence for multiple independent gains or losses across the mammalian phylogeny.

**Figure 3.**
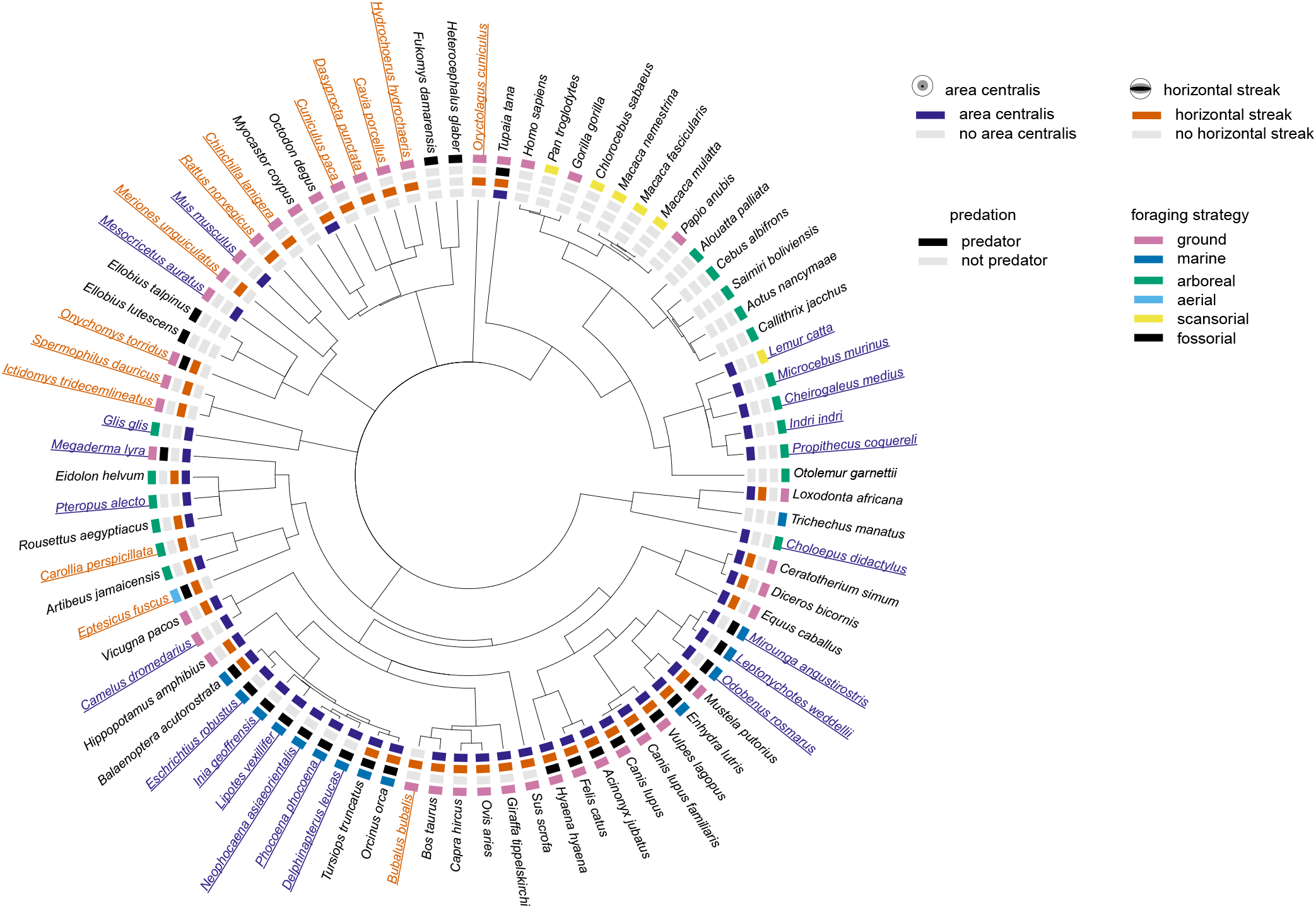
Retinal specializations overlayed on a mammalian cladogram, based on the tree topology from (Genereux et al., 2020). Colored boxes indicate the retinal specializations and ecological traits for each species. Species in purple have only an area centralis, and species in orange have only a horizontal streak.

We tested for an association between predation and the presence of an area centralis across the mammal phylogeny and did not find support for this hypothesis (LRT = 4.043; BH-corrected P = 0.185; Table 1). This was true for both species with any presence of an area centralis and only an area centralis (Table 1). We also tested if the area centralis was associated with predators that forage in three-dimensional environments (arboreal and marine), where depth perception may be particularly important. We observed a trend towards these species having an area centralis, but it was not significant after multiple test correction (LRT = 6.511; BH-corrected P = 0.096; Table 1). As a proxy for horizon-dominated habitats, we tested for an association between ground foraging and the presence of a horizontal streak and found a significant relationship between these traits (LRT = 15.938; BH-corrected P=0.003; Table 1). This relationship was no longer significant when we narrowed the foreground set to species with only a horizontal streak. The horizontal streak may be particularly common in species in horizon-dominated habitats that experience high pre-dation, because high visual acuity across the center plane may help detect predators across a wider field of view. We tested if the horizontal streak is associated with ground foraging herbivores with body mass less than 100kg, as a proxy for prey species. These likely prey species were more likely to have only a horizontal streak (LRT = 9.238; BH-corrected P = 0.049; Table 1). We also asked if species in unobstructed, horizon-dominated environments were more likely to have horizontal streaks, a prediction known as the terrain hypothesis (Hughes, 1977). We considered species to be in “unobstructed” habitats if they primarily reside in grassland, savannah, or tundra biomes (Myers et al., 2024) and excluded small rodents that are unlikely to experience unobstructed fields of view even in these open habitat types (Supplemental Table S1). These species trended towards having a horizontal streak, but the relationship was not significant after multiple test correction (LRT = 6.540; BH-corrected P = 0.096; Table 1). Our results suggest that some retinal specializations may be evolving in response to ecological conditions. In particular, the horizontal streak appears to be associated with ground foraging.

**Table 1.**
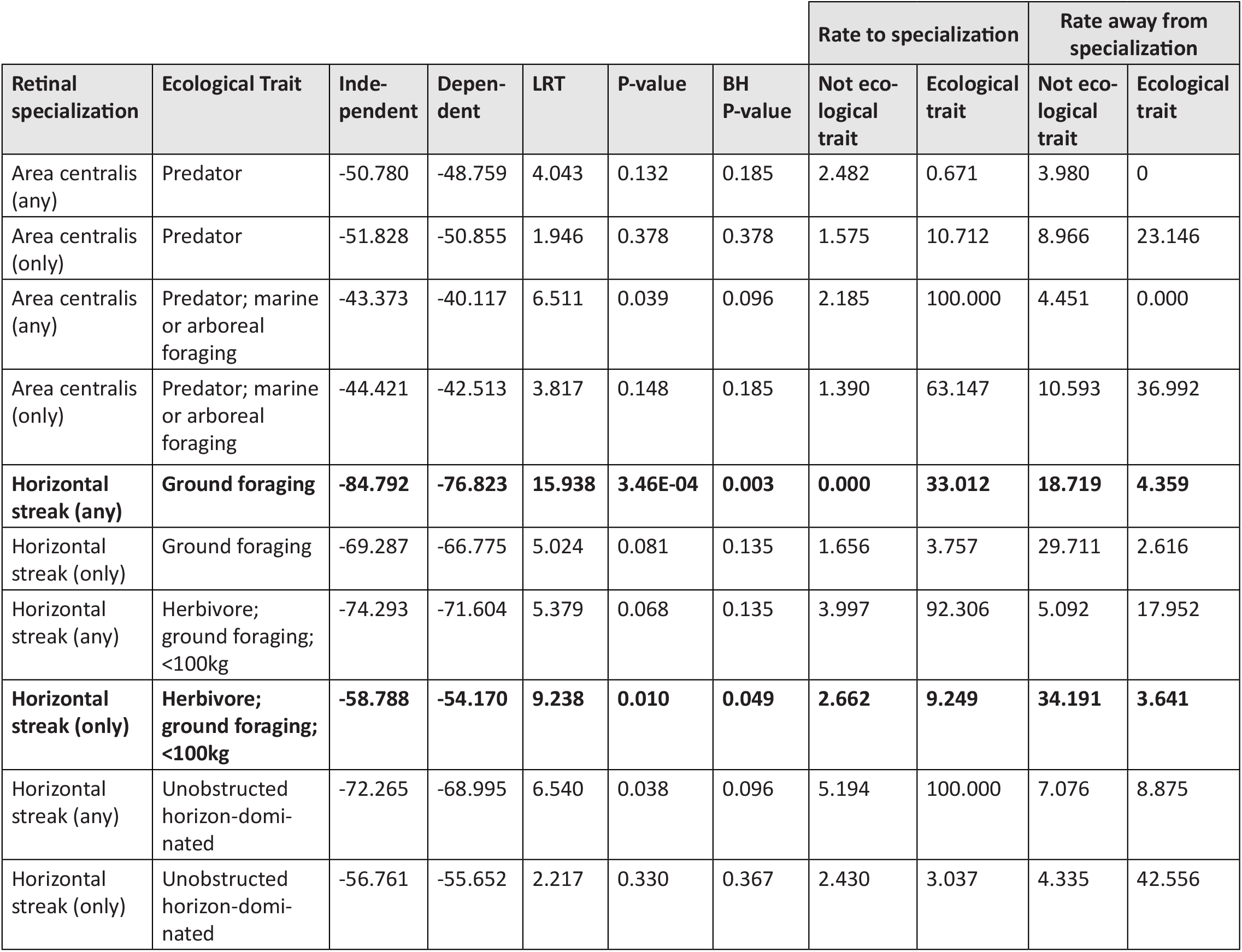
Results from hypotheses tests comparing retinal specializations to ecological traits. The independent column gives the log likelihood of the null model in which the two traits are evolving indepently across the phylogeny. The dependent column gives the log likelihood value of the model in which the evolution of one trait depends on the other. LRT is the likelihood ratio test value, the P-values were determined using a chi-squared test with two degrees of freedom, and BH P-values were corrected for multiple tests using the Benjamini-Hochberg method. The last four columns show the rates to or away from the given retinal specialization given the presence or absence of the ecological trait. Rows in bold are significant after multiple test correction.

| Retinal specialization | Ecological Trait | Independent | Dependent | LRT | P-value | BH P-value | Rate to specialization |  | Rate away from specialization |  |
| --- | --- | --- | --- | --- | --- | --- | --- | --- | --- | --- |
|  |  |  |  |  |  |  | Not ecological trait | Ecological trait | Not ecological trait | Ecological trait |
| Area centralis (any) | Predator | -50.780 | -48.759 | 4.043 | 0.132 | 0.185 | 2.482 | 0.671 | 3.980 | 0 |
| Area centralis (only) | Predator | -51.828 | -50.855 | 1.946 | 0.378 | 0.378 | 1.575 | 10.712 | 8.966 | 23.146 |
| Area centralis (any) | Predator; marine or arboreal foraging | -43.373 | -40.117 | 6.511 | 0.039 | 0.096 | 2.185 | 100.000 | 4.451 | 0.000 |
| Area centralis (only) | Predator; marine or arboreal foraging | -44.421 | -42.513 | 3.817 | 0.148 | 0.185 | 1.390 | 63.147 | 10.593 | 36.992 |
| **Horizontal streak (any)** | **Ground foraging** | **-84.792** | **-76.823** | **15.938** | **3.46E-04** | **0.003** | **0.000** | **33.012** | **18.719** | **4.359** |
| Horizontal streak (only) | Ground foraging | -69.287 | -66.775 | 5.024 | 0.081 | 0.135 | 1.656 | 3.757 | 29.711 | 2.616 |
| Horizontal streak (any) | Herbivore; ground foraging; <100kg | -74.293 | -71.604 | 5.379 | 0.068 | 0.135 | 3.997 | 92.306 | 5.092 | 17.952 |
| **Horizontal streak (only)** | **Herbivore; ground foraging; <100kg** | **-58.788** | **-54.170** | **9.238** | **0.010** | **0.049** | **2.662** | **9.249** | **34.191** | **3.641** |
| Horizontal streak (any) | Unobstructed horizon-dominated | -72.265 | -68.995 | 6.540 | 0.038 | 0.096 | 5.194 | 100.000 | 7.076 | 8.875 |
| Horizontal streak (only) | Unobstructed horizon-dominated | -56.761 | -55.652 | 2.217 | 0.330 | 0.367 | 2.430 | 3.037 | 4.335 | 42.556 |

### There is a strong association between orbit convergence and retinal specializations

We then asked if there is a relationship between orbit convergence and retinal specializations. Both the horizontal streak and more side-facing eyes are thought to facilitate panoramic vision in open, horizon-dominated environments, and therefore we predicted that they would tend to occur together (Hughes, 1977). Indeed, we found that orbit convergence was significantly lower in species that had only a horizontal streak compared to species that had only a fovea or area centralis (PGLS P = 0.0398; Figure 4A). We also tested this relationship using a simulation-based phylogenetic ANOVA (Garland et al., 1993) and again saw significantly lower orbit convergence for species with only a horizontal streak (P = 0.0195).

**Figure 4.**
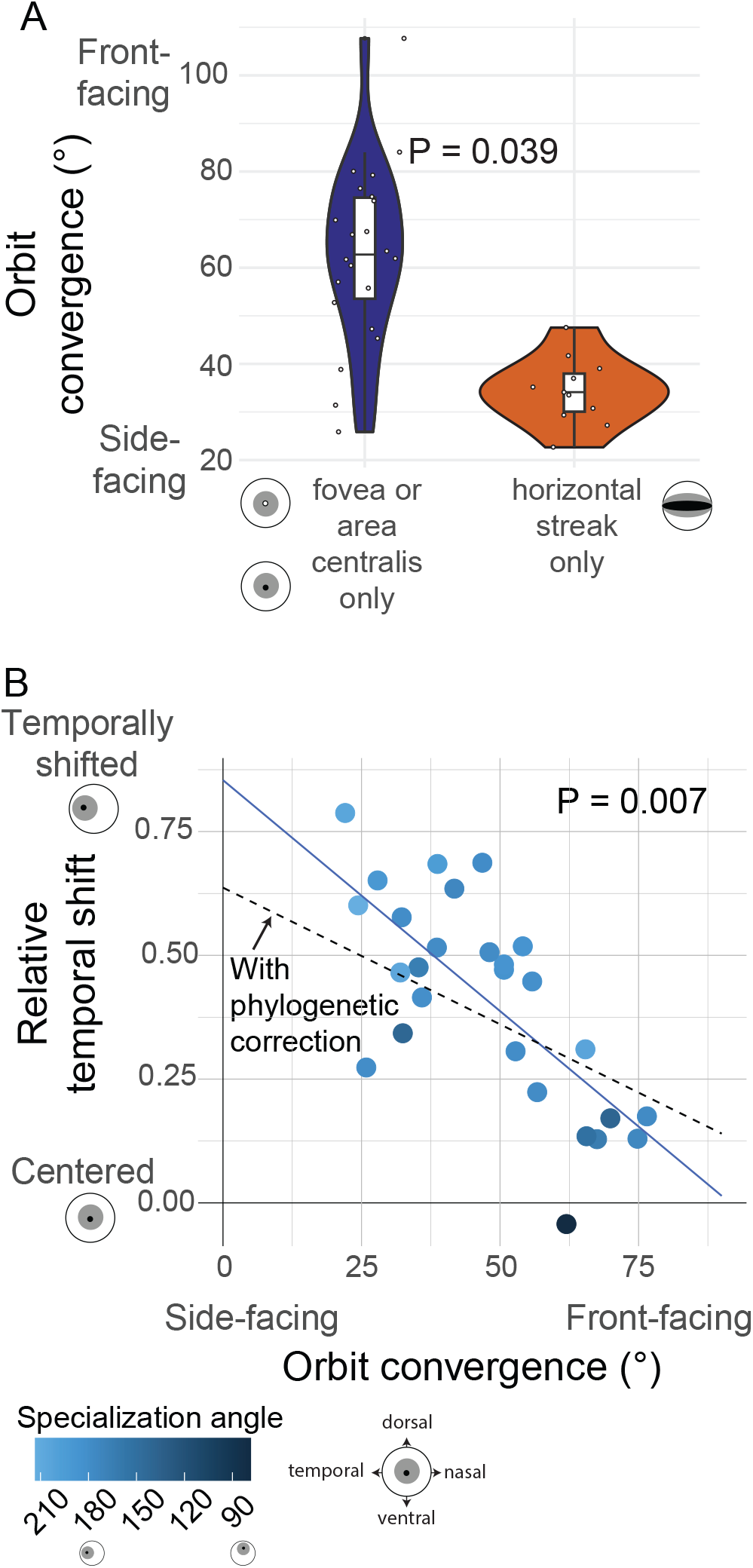
The relationship between retinal specializations and orbit convergence. (A) Violin plot showing the relationship between orbit convergence and presence of different retinal specialization types. Each point represents a species, and thick horizontal lines within the violin plot represent the median orbit convergence angle for species with the given retinal specialization. (B) Scatter plot showing the negative correlation between orbit convergence and relative temporal distance of the fovea or area centralis from the center of the retina. Points are colored based on the angle of the retinal specialization relative to the horizontal plane of the retina. The orange line represents the slope and intercept of the phylogenetic generalized least squares analysis, and the blue line shows the regression without phylogenetic correction. P-values are based on phylogenetic generalized least squares with body mass as a covariate.

Previous studies have observed that species with lower orbit convergence tend to have temporally-shifted retinal specializations, which would allow for higher acuity in the binocular field, the region in front of an animal where the visual fields of both eyes overlap [Figure 1C; (Collin, 1999; Moore et al., 2012)]. Essentially, in these cases the lateral shift serves to redirect the high acuity area so that it faces more forward with respect to the head, despite the side-facing orientation of the eyes themselves. We asked if the relative temporal distance of the area centralis or fovea from the center of the retina was correlated with orbit convergence and found strong support for an inverse relationship between these measurements (PGLS P = 0.0071; Figure 4B). That is, species with more laterally placed eyes tend to have more temporally shifted retinal specializations, as predicted. The regression line nearly intercepts the x-axis at 90 degrees and a temporal shift of zero, which means species with eyes facing directly forward would have an almost centered retinal specialization. The y-intercept is close to 1, which would mean that species with eyes facing directly sideways would have retinal specializations shifted completely to the temporal side of the retina. This suggests that most species with a high acuity retinal specialization have a region of highest acuity that projects almost directly forwards, regardless of the position of the eyes in the head.

## Discussion

It is still unclear how selective pressures influence many ocular traits, or what evolutionary forces underlie divergence in these traits across species. Ecological correlates of ocu-lar traits can provide insight into their potential roles as visual adaptations (Caves et al., 2017; Potier et al., 2017; Baker & Venditti, 2019; Cantlay et al., 2023; Potier et al., 2023; Caves et al., 2024). Additionally, multiple traits involved in vision are predicted to evolve in response to similar ecological conditions, but the way these traits interact with each other may also shape their evolutionary patterns (Walls, 1942; Hughes, 1977; Collin, 1999). We tested hypotheses regarding the ecological selective pressures acting on orbit convergence and high acuity retinal specializations and found mixed evidence supporting these hypotheses. However, we did find strong evidence for correlations among these traits across the mammal phylogeny, suggesting that functional interactions between ocular traits may play an important role in their evolution.

### Ecological correlates of ocular traits

Phylogenetic comparative methods can give insight into adaptive evolution, especially for traits evolving over longer evolutionary timescales that cannot be directly tested for selection within populations (Harmon, 2019). As genome sequencing has become more accessible and affordable, phylogenetic inference has improved, generating well-resolved phylogenetic trees consisting of many species across long evolutionary timescales (Shen et al., 2018; Upham et al., 2019; Feng et al., 2020; Genereux et al., 2020; Suvorov et al., 2022). While the importance of controlling for shared evolutionary history is well-established (Felsenstein, 1985; Harmon, 2019), many older hypotheses about the relationships among traits and ecological conditions have not been revisited using current phylogenies and phylogenetic comparative methods. Testing these hypotheses with modern phylogenetic methods and resources can reveal new insights about trait evolution and putative ecological selective pressures acting on traits (Baker et al., 2016; Davis et al., 2016; Baker & Venditti, 2019; Benun Sutton & Wilson, 2019; Harmon, 2019; Jarvis & Marshall, 2023).

We tested several long-standing hypotheses regarding ecological selective pressures acting on visual traits. Surprisingly, we did not find support for most of these hypotheses in mammals. After controlling for phylogeny, we found that orbit convergence was not associated with diet, activity pattern, or foraging substrate (Figure 2). This contrasts with previous work that did not account for shared evolutionary history (Heesy, 2007), but is consistent with findings in carnivores and marsupials (Pilatti & Astúa, 2016; Casares-Hidalgo et al., 2019). Phylogenetic relationships and ecological correlates thought to influence orbit convergence may be confounded in mammals, because high orbit convergence seems to be concentrated in one or a few mammalian clades with similar relevant ecological traits. For example, carnivores tend to have high orbit convergence associated with predation, but they are also a monophyletic clade and may have similar orbit convergence due to their shared evolutionary history (Casares-Hidalgo et al., 2019). These results also differ from findings in birds, which show strong correlations between the degree of binocular overlap and ecological traits such as diet and activity pattern based on phylogenetic comparative models (Cantlay et al., 2023; Potier et al., 2023). Birds may be even more visually-oriented than mammals and rely less on other sensory systems such as smell, so their ocular traits may evolve more rapidly in response to ecological selective pressures on vision (Caves et al., 2024). Although we did not find statistical support for relationships between ecology and orbit convergence, these hypothesized ecological selective pressures may still act on orbit convergence. However, orbit convergence evolves relatively slowly in mammals (Figure 2), and therefore there may not be enough independent shifts in orbit convergence across mammals to detect these associations in a phylogenetic framework.

We found some evidence for associations between retinal specializations and ecology (Figure 3). There was a weak association between the area centralis and predation in three dimensional environments. We defined predation based on diet composition because these data were widely available across species in our dataset. However, different hunting strategies may impose different selective pressures on vision, such as ambush versus active predation (Banks et al., 2015). The area centralis and other ocular traits may evolve in response to more specific selective pressures imposed by different types of predation, but we did not have behavioral data at this level of detail to make comparisons across species. We found a significant association between the horizontal streak and ground foraging, supporting the hypothesis that horizon-dominated habitats may select for the presence of a horizontal streak. We also found that species with only a horizontal streak were more likely to be small, herbivorous ground foragers. There was a weak association between ground foraging species in unobstructed habitats and the presence of the horizontal streak. However, this was not significant, and the strongest relationship we observed was with ground foraging. It may be that ground foraging alone is enough to select for the presence of a horizontal streak, regardless of how open the habitat is or the diet of a species. Previous studies have found variation in retinal specializations associated with foraging terrain complexity in artiodactyls (Schiviz et al., 2008) and marsupials (Navarro-Sempere et al., 2018). Our work shows that these patterns persist across deeper evolutionary time scales and across species with greater variation in habitats and foraging strategies.

### Evolutionary relationships between ocular traits

The strongest relationships we observed were those between orbit convergence and retinal specializations. Orbit convergence was significantly lower in species with only a horizontal streak compared to those with a fovea or area centralis (Figure 4A), which may reflect similar selective pressures for panoramic vision and a wide field of view acting on both orbit convergence and patterns of retinal ganglion cell density. We also found a strong negative correlation between the temporal shift of the high acuity specialization from the center of the retina and orbit convergence, meaning that the high acuity area shifts temporally in species with side-facing eyes (Figure 4B). Previous studies predicted this pattern, as it would provide the highest visual acuity in front of an animal even if the eyes are facing sideways (Figure 1C)(Collin, 1999; Moore et al., 2012). Some species were known to support this prediction (Hughes, 1977; Collin, 1999), but we provide, to our knowledge, the first quantitative support for this relationship in a phylogenetic framework.

Orbit convergence shows strong phylogenetic signal and evolves relatively slowly in mammals, but in contrast, retinal specializations have likely been gained and lost multiple times in mammals (Figure 2; Figure 3). Orbit morphology is likely constrained by other aspects of cranial morphology (Ross, 1995; Cox, 2008; Finarelli & Goswami, 2009; Pilatti & Astúa, 2016). While ecological selective pressures on vision may play some role in the evolution of orbit convergence, they may be difficult to detect among other factors influencing the evolution of this trait, particularly morphological constraints and the influence of shared evolutionary history (Casares-Hidalgo et al., 2019). In contrast, the arrangement of ganglion cells within the retina is likely subject to fewer constraints, and therefore may evolve to compensate for other ocular traits such as orbit convergence.

Retinal specializations are usually binned into categories due to historical precedent and available methodologies (Hughes, 1977). However, patterns of retinal ganglion cell density are often more complex than these categories, and many species have specializations that fit multiple categories (Figure 3; Supplemental Table S1). Considering quantitative parameters of retinal specializations, such as changes in ganglion cell density across the retina, provides a more complete understanding of the evolution of retinal specializations (Schiviz et al., 2008; Moore et al., 2012). Indeed, the strongest trend we observed was the relationship between the temporal shift of the specialization and orbit convergence, which required the quantitative measurement of the relative temporal distance of the specialization. Different studies have phenotyped and reported retinal specializations using different approaches, limiting the ability to test evolutionary hypotheses about quantitative measures of retinal ganglion cell density (Moore et al., 2012). Future work on retinal specializations should include detailed retinal topography maps as well as raw data on ganglion cell density across the retina, allowing comparative work to test hypotheses about the evolution of quantitative retinal specialization traits (Moore et al., 2012). In this study, we used categorical and quantitative data on retinal specializations to show that their occurrence and position within the retina likely evolved both in response to ecological selective pressures and to compensate for other ocular traits. Vision, like many sensory processes, requires the coordination of multiple traits across molecular, cellular, and morphological levels. Testing how these traits evolve in relation to one another in the context of visual ecology provides a more complete understanding of the evolution of sensory systems.

## Author contributions

E.E.K.K. performed the literature searches and data analyses with support from N.L.C. E.E.K.K. and N.L.C. funded this work, collected data on orbit convergence, and wrote the manuscript.

## Acknowledgements

We thank members of the Clark lab, Esteban Fernandez-Juricic, and Bret Moore for helpful feedback on this work. Dwon Jordana, Courtney Charlesworth, and Alex Preble assisted with orbit convergence measurements. Thank you to John Wible at the Carnegie Museum of Natural History for providing specimens and Shawn Artman at the University of Pittsburgh machine shop for building the custom dihedral goniometers. We thank Matt Ravosa for providing information about the construction and use of the goniometer.

## Funding

This research was supported in part by the University of Pittsburgh Center for Research Computing, RRID:SCR_022735, through the resources provided. Specifically, this work used the HTC cluster, which is supported by NIH award number S10OD028483. This work was supported by a National Science Foundation Postdoctoral Fellowship in Biology to E.E.K.K. under Grant No. 2305797. Any opinions, findings, and conclusions or recommendations expressed in this material are those of the author(s) and do not necessarily reflect the views of the National Science Foundation or the National Institutes of Health.

## Conflict of Interest Statement

The authors declare no conflicts of interest.

## Notes

### Competing Interest Statement

The authors have declared no competing interest.

