## supplemental material for "Mammalian retinal specializations for high acuity vision evolve in response to both foraging strategies and morphological constraints"

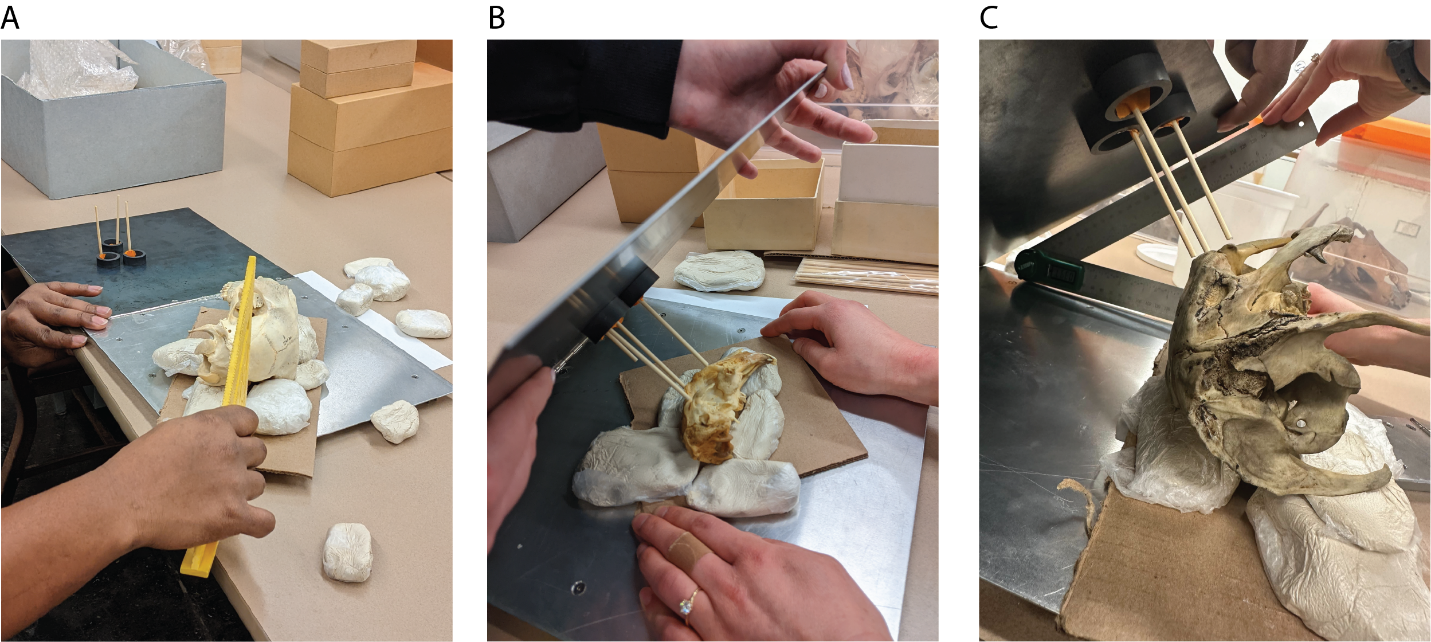

**Figure S1:** Dihedral goniometer. (A) The specimen is balanced on its side using clay, and positions such that the prostion, inion, and basion points that define the sagittal plane are an equal distance from the base of the goniometer. This ensures that the sagittal plane is parallel with the base of the goniometer. (B) The top of the goniometer is folded down and the position of the skull and magnetic pins adjusted until all three pins touch three points on the orbital plane. (C) The angle between the top and base of the goniometer is measured with a digital protractor. This is the orbit convergence angle. Method adapted from (Ross, 1995). Photos by Emily Kopania and Courtney Charlesworth.

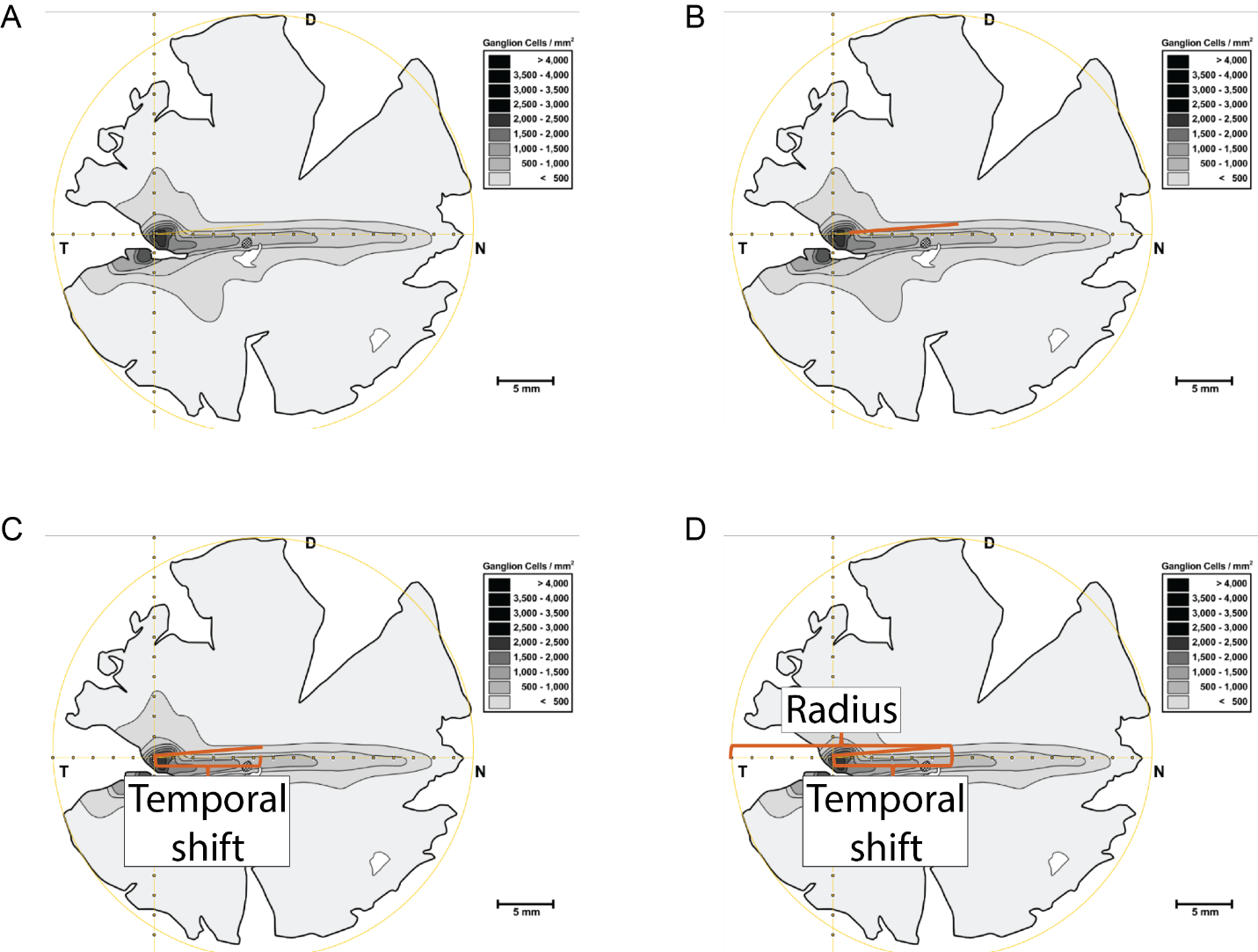

**Figure S2:** Relative temporal shift. (A) The extreme points of the retinal topographic map were manually marked on the dorsal, nasal, ventral, and temporal sides. Built-in ImageJ functions were used to draw a minimum bounding circle (yellow). The center of the high acuity retinal specialization was also manually marked (intersection of yellow lines). (B) Built-in ImageJ functions were used to measure the distance from the center of the minimum bounding circle, indicating the center of the retina, and the center of the retinal specialization (orange line). (C) Built-in ImageJ functions were used to calculate the angle between the retinal specialization and the horizontal line intersecting the center of the retina, and trigonometry functions were used to calculate the temporal shift, or the distance of the specialization away from the center of the retina along the horizontal axis (bracket). (D) The temporal shift was divided by the radius (large bracket) to calculate the relative temporal shift, normalized for different sizes of retinas and different units of measurement. Method adapted from (Moore et al., 2012). Retinal topography map image from (Calderone et al., 2003). Scripts for ImageJ macros and trigonometry functions available at <https://github.com/ekopania/mammal_retinas>.

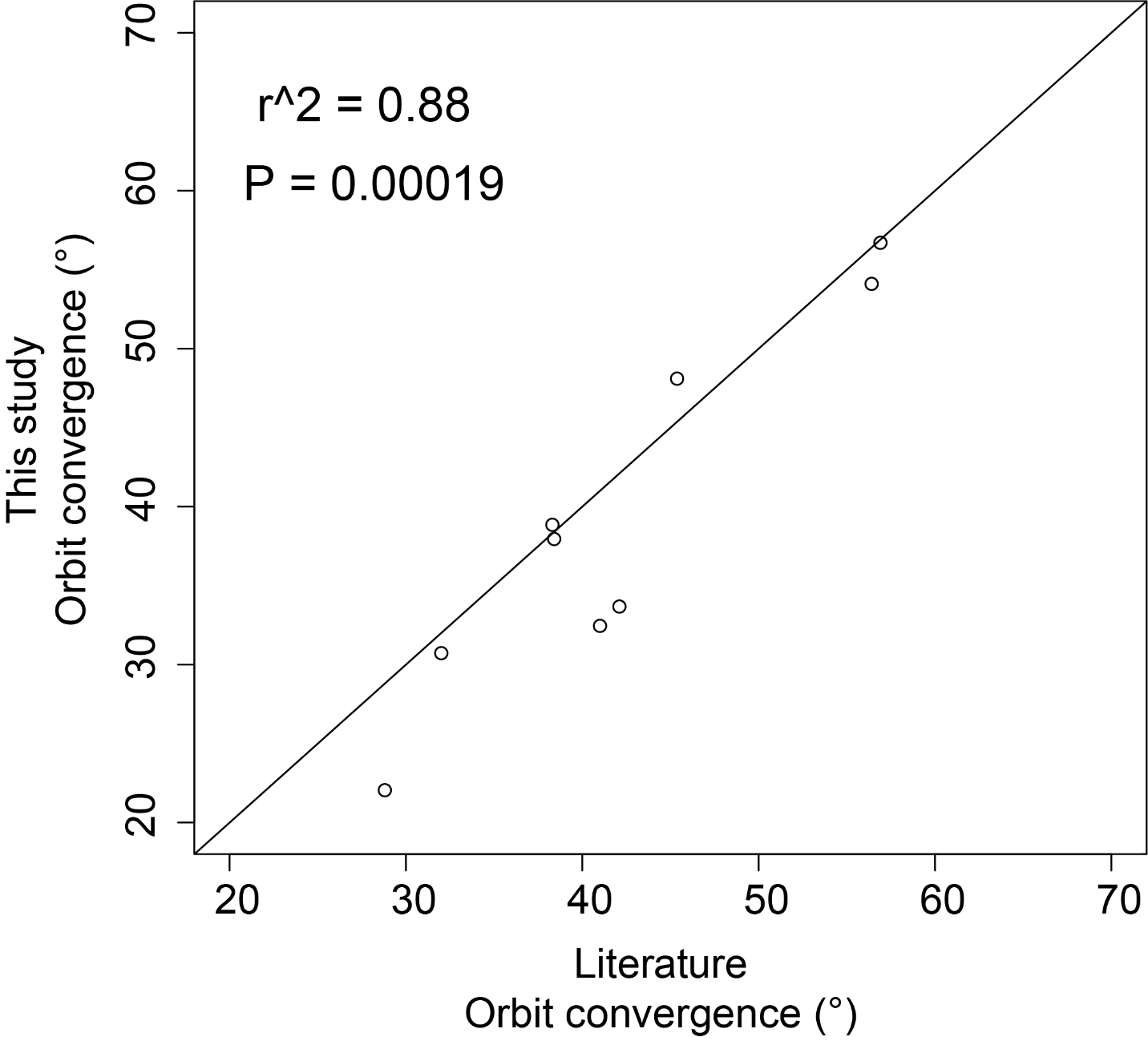

**Figure S3:** Orbit convergence measurements taken for this study compared to published orbit convergence measurements from the literature for the same species. Each point represents a species, with its measurement from this study on the y-axis and its measurement from the literature on the x-axis. R^2^ and p-value based on a Pearson’s correlation test.

**Supplemental References, including sources from Supplemental Table S1**

Bringmann, A., Syrbe, S., Görner, K., Kacza, J., Francke, M., Wiedemann, P., & Reichenbach, A. (2018). The primate fovea: Structure, function and development. *Prog. Retin. Eye Res., 66*, 49-84. doi:<https://doi.org/10.1016/j.preteyeres.2018.03.006>

Calderone, J. B., Reese, B. E., & Jacobs, G. H. (2003). Topography of photoreceptors and retinal ganglion cells in the spotted hyena (*Crocuta crocuta*). *Brain Behav Evol, 62*(4), 182-192. doi:10.1159/000073270

Cechetto, C., de Busserolles, F., Jakobsen, L., & Warrant, E. J. (2020). Retinal ganglion cell topography and spatial resolving power in echolocating and non-echolocating bats. *Brain Behav. Evol., 95*(2), 58-68. doi:10.1159/000508863

Coimbra, J. P., Bertelsen, M. F., & Manger, P. R. (2017a). Retinal ganglion cell topography and spatial resolving power in the river hippopotamus (*Hippopotamus amphibius*). *J. Comp. Neurol., 525*(11), 2499-2513. doi:<https://doi.org/10.1002/cne.24179>

Coimbra, J. P., Hart, N. S., Collin, S. P., & Manger, P. R. (2013). Scene from above: Retinal ganglion cell topography and spatial resolving power in the giraffe (*Giraffa camelopardalis*). *J. Comp. Neurol., 521*(9), 2042-2057. doi:<https://doi.org/10.1002/cne.23271>

Coimbra, J. P., & Manger, P. R. (2017). Retinal ganglion cell topography and spatial resolving power in the white rhinoceros (*Ceratotherium simum*). *J. Comp. Neurol., 525*(11), 2484-2498. doi:<https://doi.org/10.1002/cne.24136>

Coimbra, J. P., Pettigrew, J. D., Kaswera-Kyamakya, C., Gilissen, E., Collin, S. P., & Manger, P. R. (2017b). Retinal ganglion cell topography and spatial resolving power in African megachiropterans: Influence of roosting microhabitat and foraging. *J. Comp. Neurol., 525*(1), 186-203. doi:<https://doi.org/10.1002/cne.24055>

Curcio, C. A., & Allen, K. A. (1990). Topography of ganglion cells in human retina. *J. Comp. Neurol., 300*(1), 5-25. doi:<https://doi.org/10.1002/cne.903000103>

Debruyn, E. J., III. (1983). *The organization and central terminations of retinal ganglion cells in the tree shrew (Tupaia glis).* (Ph.D. Doctoral dissertation), Vanderbilt University, United States -- Tennessee. Retrieved from <http://pitt.idm.oclc.org/login?url=https://www.proquest.com/dissertations-theses/organization-central-terminations-retinal/docview/303195615/se-2?accountid=14709> ProQuest Dissertations & Theses Global database. (8317333)

Dkhissi-Benyahya, O., Szel, A., Degrip, W. J., & Cooper, H. M. (2001). Short and mid-wavelength cone distribution in a nocturnal Strepsirrhine primate (*Microcebus murinus*). *J. Comp. Neurol., 438*(4), 490-504. doi:<https://doi.org/10.1002/cne.1330>

Do-Nascimento, J. L., Do-Nascimento, R. S., Damasceno, B. A., & Silveira, L. C. (1991). The neurons of the retinal ganglion cell layer of the guinea pig: quantitative analysis of their distribution and size. *Braz J Med Biol Res, 24*(2), 199-214.

Fischer, Q. S., & Kirby, M. A. (1991). Number and Distribution of Retinal Ganglion Cells in Anubis Baboons (*Papio anubis*). *Brain Behav Evol, 37*(4), 189-203. doi:10.1159/000114358

Gao, A., & Zhou, K. (1987). On the retinal ganglion cells of neophocaena and lipotes. *Acta Zool Sin, 33*(4), 316-322.

Garciá, M., Ruiz-Ederra, J., Hernández-Barbáchano, H., & Vecino, E. (2005). Topography of pig retinal ganglion cells. *J. Comp. Neurol., 486*(4), 361-372. doi:<https://doi.org/10.1002/cne.20516>

Gilmore, D. P., Da-Costa, C. P., & Duarte, D. P. (2000). An update on the physiology of two- and three-toed sloths. *Braz J Med Biol Res, 33*(2), 129-146. doi:10.1590/s0100-879x2000000200001

Gonzalez-Soriano, J., Mayayo-Vicente, S., Martinez-Sainz, P., Contreras-Rodriguez, J., & Rodriguez-Veiga, E. (1997). A quantitative study of ganglion cells in the goat retina. *Anat. Histol. Embryol., 26*(1), 39-44. doi:<https://doi.org/10.1111/j.1439-0264.1997.tb00101.x>

Guo, X., & Sugita, S. (2000). Topography of ganglion cells in the retina of the horse. *J. Vet. Med. Sci., 62*(11), 1145-1150. doi:10.1292/jvms.62.1145

Harman, A., Dann, J., Ahmat, A., Macuda, T., Johnston, K., & Timneyb, B. (2001). The retinal ganglion cell layer and visual acuity of the camel. *Brain Behav. Evol., 58*(1), 15-27. doi:10.1159/000047258

Hebel, R. (1976). Distribution of retinal ganglion cells in five mammalian species (pig, sheep, ox, horse, dog). *Anat. Embryol. (Berl.), 150*(1), 45-51. doi:10.1007/BF00346285

Heffner, R. S., & Heffner, H. E. (1992). Visual factors in sound localization in mammals. *J. Comp. Neurol., 317*(3), 219-232. doi:<https://doi.org/10.1002/cne.903170302>

Heffner, R. S., Heffner, H. E., Kearns, D., Vogel, J., & Koay, G. (1994). Sound localization in chinchillas. I: Left/right discriminations. *Hearing Res., 80*(2), 247-257. doi:<https://doi.org/10.1016/0378-5955(94)90116-3>

Heffner, R. S., Koay, G., & Heffner, H. E. (2001). Sound localization in a new-world frugivorous bat, *Artibeus jamaicensis*: Acuity, use of binaural cues, and relationship to vision. *The Journal of the Acoustical Society of America, 109*(1), 412-421. doi:10.1121/1.1329620

Heffner, R. S., Koay, G., & Heffner, H. E. (2007). Sound-localization acuity and its relation to vision in large and small fruit-eating bats: I. Echolocating species, *Phyllostomus hastatus* and *Carollia perspicillata*. *Hearing Res., 234*(1), 1-9. doi:<https://doi.org/10.1016/j.heares.2007.06.001>

Heffner, R. S., Koay, G., & Heffner, H. E. (2008). Sound localization acuity and its relation to vision in large and small fruit-eating bats: II. Non-echolocating species, *Eidolon helvum* and *Cynopterus brachyotis*. *Hearing Res., 241*(1), 80-86. doi:<https://doi.org/10.1016/j.heares.2008.05.001>

Henderson, Z. (1985). Distribution of ganglion cells in the retina of adult pigmented ferret. *Brain Res., 358*(1), 221-228. doi:<https://doi.org/10.1016/0006-8993(85)90966-7>

Herbin, M., Boire, D., & Ptito, M. (1997). Size and distribution of retinal ganglion cells in the St. Kitts green monkey (*Cercopithecus aethiops sabeus)*. *J. Comp. Neurol., 383*(4), 459-472. doi:[https://doi.org/10.1002/(SICI)1096-9861(19970714)383:4<459::AID-CNE5>3.0.CO;2-1](https://doi.org/10.1002/(SICI)1096-9861(19970714)383:4%3c459::AID-CNE5%3e3.0.CO;2-1)

Herbin, M., Repérant, J., & Cooper, H. M. (1994). Visual system of the fossorial mole-lemmings, *Ellobius talpinus* and *Ellobius lutescens*. *J. Comp. Neurol., 346*(2), 253-275. doi:<https://doi.org/10.1002/cne.903460206>

Heukamp, A. S., Warwick, R. A., & Rivlin-Etzion, M. (2020). Topographic variations in retinal encoding of visual space. *Annual Review of Vision Science, 6*(1), 237-259. doi:10.1146/annurev-vision-121219-081831

Hughes, A. (1975). A quantitative analysis of the cat retinal ganglion cell topography. *J. Comp. Neurol., 163*(1), 107-128. doi:<https://doi.org/10.1002/cne.901630107>

Hughes, A. (1977). The topography of vision in mammals of contrasting life style: Comparative optics and retinal organisation. In F. Crescitelli, C. A. Dvorak, D. J. Eder, A. M. Granda, D. Hamasaki, K. Holmberg, A. Hughes, N. A. Locket, W. N. McFarland, D. B. Meyer, W. R. A. Muntz, F. W. Munz, E. C. Olson, R. W. Reyer, & F. Crescitelli (Eds.), *The Visual System in Vertebrates* (pp. 613-756). Berlin, Heidelberg: Springer Berlin Heidelberg.

Kassab, A., & Sugita, S. (2000). Study of ganglion cell topography of the retina in buffaloes (*Bos bubalis*). *日本畜産学会報, 71*(6), 600-608. doi:10.2508/chikusan.71.600

Knapp, S., McCulley, J. P., Alvarado, T. P., & Hogan, R. N. (2007). Comparative ocular anatomy of the western lowland gorilla. *Vet. Ophthalmol., 10*(6), 357-362. doi:<https://doi.org/10.1111/j.1463-5224.2007.00568.x>

Koay, G., Kearns, D., Heffner, H. E., & Heffner, R. S. (1998). Passive sound-localization ability of the big brown bat (*Eptesicus fuscus*). *Hearing Res., 119*(1), 37-48. doi:<https://doi.org/10.1016/S0378-5955(98)00037-9>

Long, K. O., & Fisher, S. K. (1983). The distributions of photoreceptors and ganglion cells in the California ground squirrel, *Spermophilus beecheyi*. *J. Comp. Neurol., 221*(3), 329-340. doi:<https://doi.org/10.1002/cne.902210308>

Malkemper, E. P., & Peichl, L. (2018). Retinal photoreceptor and ganglion cell types and topographies in the red fox (*Vulpes vulpes*) and Arctic fox (*Vulpes lagopus*). *J. Comp. Neurol., 526*(13), 2078-2098. doi:<https://doi.org/10.1002/cne.24493>

Mass, A. M. (2001). Visual field organization and retinal resolution in the beluga whale *Delphinapterus leucas* (Pallas). *Dokl. Biol. Sci., 381*(1), 555-558. doi:10.1023/A:1013326521559

Mass, A. M., Ketten, D. R., Odell, D. K., & Supin, A. Y. (2012). Ganglion cell distribution and retinal resolution in the Florida manatee, *Trichechus manatus latirostris*. *The Anatomical Record, 295*(1), 177-186. doi:<https://doi.org/10.1002/ar.21470>

Mass, A. M., & Supin, A. Y. (1989). Distribution of ganglion cells in the retina of an Amazon River dolphin. *Aquat. Mamm., 15*(2), 49-56.

Mass, A. M., & Supin, A. Y. (2007). Adaptive features of aquatic mammals' eye. *The Anatomical Record, 290*(6), 701-715. doi:<https://doi.org/10.1002/ar.20529>

Mass, A. M., & Supin, A. Y. (2017). Estimates of underwater and aerial visual acuity in the European beaver *Castor fiber* L. based on morphological data. *Dokl. Biol. Sci., 473*(1), 35-38. doi:10.1134/S0012496617020065

Mass, A. M., Supin, A. Y., Abramov, A. V., Mukhametov, L. M., & Rozanova, E. I. (2013). Ocular anatomy, ganglion cell distribution and retinal resolution of a killer whale (*Orcinus orca*). *Brain Behav. Evol., 81*(1), 1-11. doi:10.1159/000341949

Mass, A. M., Supin, A. Y., & Severtsov, A. N. (1986). Topographic distribution of sizes and density of ganglion cells in the retina of a porpoise, *Phocoena phocoena*. *Aquat. Mamm., 12*(3), 95-102.

Mass, A. M., & Ya, A. (1997). Ocular anatomy, retinal ganglion cell distribution, and visual resolution in the gray whale, *Eschrichtius gibbosus*. *Aquat. Mamm., 23*, 17–28.

Mass, A. M., & Ya. Supin, A. (2000). Ganglion cells density and retinal resolution in the sea otter, *Enhydra lutris*. *Brain Behav. Evol., 55*(3), 111-119. doi:10.1159/000006646

Miyazaki, T., Naritsuka, Y., Yagami, M., Kobayashi, S., & Kawamura, K. (2022). Anatomy and histology of the eye of the nutria *Myocastor coypus*: Evidence of adaptation to a semi-aquatic life. *Zool Stud, 61*, e18. doi:10.6620/zs.2022.61-18

Moore, B. A., Kamilar, J. M., Collin, S. P., Bininda-Emonds, O. R. P., Dominy, N. J., Hall, M. I., Heesy, C. P., Johnsen, S., Lisney, T. J., Loew, E. R., Moritz, G., Nava, S. S., Warrant, E., Yopak, K. E., & Fernández-Juricic, E. (2012). A novel method for comparative analysis of retinal specialization traits from topographic maps. *J Vision, 12*(12), 13-13. doi:10.1167/12.12.13

Muniz, J. A. P. C., de Athaide, L. M., Gomes, B. D., Finlay, B. L., & Silveira, L. C. d. L. (2015). Ganglion cell and displaced amacrine cell density distribution in the retina of the howler monkey (*Alouatta caraya*). *PLoS One, 9*(12), e115291. doi:10.1371/journal.pone.0115291

Murayama, T., Somiya, H., Aoki, I., & Ishii, T. (1992). The distribution of ganglion cells in the retina and visual acuity of minke whale. *Nippon Suisan Gakkaishi, 58*(6), 1057-1061.

Němec, P., Cveková, P., Benada, O., Wielkopolska, E., Olkowicz, S., Turlejski, K., Burda, H., Bennett, N. C., & Peichl, L. (2008). The visual system in subterranean African mole-rats (Rodentia, Bathyergidae): Retina, subcortical visual nuclei and primary visual cortex. *Brain Res. Bull., 75*(2), 356-364. doi:<https://doi.org/10.1016/j.brainresbull.2007.10.055>

Packer, O., Hendrickson, A. E., & Curcio, C. A. (1989). Photoreceptor topography of the retina in the adult pigtail macaque (*Macaca nemestrina*). *J. Comp. Neurol., 288*(1), 165-183. doi:<https://doi.org/10.1002/cne.902880113>

Peichl, L. (1992). Topography of ganglion cells in the dog and wolf retina. *J. Comp. Neurol., 324*(4), 603-620. doi:<https://doi.org/10.1002/cne.903240412>

Peichl, L., Radic, T., Solovei, I., Wolfram, M., & Glösmann, M. (2022). On the retinae of Glis and Graphiurus: photoreceptor and ganglion cell populations, an absence of shortwave-sensitive cones, and some other features (Rodentia: Gliridae). *Lynx (Prague), 53*(1), 185-205. doi:<https://doi.org/10.37520/lynx.2022.013>

Perry, V. H., & Cowey, A. (1985). The ganglion cell and cone distributions in the monkey's retina: Implications for central magnification factors. *Vision Res., 25*(12), 1795-1810. doi:<https://doi.org/10.1016/0042-6989(85)90004-5>

Pettigrew, J. D., Bhagwandin, A., Haagensen, M., & Manger, P. R. (2010). Visual acuity and heterogeneities of retinal ganglion cell densities and the tapetum lucidum of the african elephant (*Loxodonta africana*). *Brain Behav. Evol., 75*(4), 251-261. doi:10.1159/000314898

Pettigrew, J. D., Dreher, B., Hopkins, C. S., McCall, M. J., & Brown, M. (1988). Peak density and distribution of ganglion cells in the retinae of microchiropteran bats: Implications for visual acuity (part 1 of 2). *Brain Behav Evol, 32*(1), 39-56. doi:10.1159/000116531

Pettigrew, J. D., & Manger, P. R. (2008). Retinal ganglion cell density of the black rhinoceros (*Diceros bicornis*): Calculating visual resolution. *Vis. Neurosci., 25*(2), 215-220. doi:10.1017/S0952523808080498

Prestrude, A. M. (1970). Sensory capacities of the chimpanzee: A review. *Psychol. Bull., 74*(1), 47-67. doi:<https://doi.org/10.1037/h0029404>

Provis, J. M., Diaz, C. M., & Dreher, B. (1998). Ontogeny of the primate fovea:a central issue in retinal development. *Prog. Neurobiol., 54*(5), 549-581. doi:<https://doi.org/10.1016/S0301-0082(97)00079-8>

Rohen, J. W., & Castenholz, A. (2008). Über die zentralisation der retina bei primaten. *Folia Primatol., 5*(1-2), 92-147. doi:10.1159/000161941

Ross, C. F. (1995). Allometric and functional influences on primate orbit orientation and the origins of the Anthropoidea. *J. Hum. Evol., 29*(3), 201-227. doi:<https://doi.org/10.1006/jhev.1995.1057>

Salinas-Navarro, M., Mayor-Torroglosa, S., Jiménez-López, M., Avilés-Trigueros, M., Holmes, T. M., Lund, R. D., Villegas-Pérez, M. P., & Vidal-Sanz, M. (2009). A computerized analysis of the entire retinal ganglion cell population and its spatial distribution in adult rats. *Vision Res., 49*(1), 115-126. doi:<https://doi.org/10.1016/j.visres.2008.09.029>

Shinozaki, A., Hosaka, Y., Imagawa, T., & Uehara, M. (2010). Topography of ganglion cells and photoreceptors in the sheep retina. *J. Comp. Neurol., 518*(12), 2305-2315. doi:<https://doi.org/10.1002/cne.22333>

Silveira, L. C. L., Perry, V. H., & Yamada, E. S. (1993). The retinal ganglion cell distribution and the representation of the visual field in area 17 of the owl monkey, *Aotus trivirgatus*. *Vis. Neurosci., 10*(5), 887-897. doi:10.1017/S095252380000609X

Silveira, L. C. L., Picanço-Diniz, C. W., & Oswaldo-Cruz, E. (1989a). Distribution and size of ganglion cells in the retinae of large Amazon rodents. *Vis. Neurosci., 2*(3), 221-235. doi:10.1017/S0952523800001140

Silveira, L. C. L., Picanço-Diniz, C. W., Sampaio, L. F. S., & Oswaldo-Cruz, E. (1989b). Retinal ganglion cell distribution in the Cebus monkey: A comparison with the cortical magnification factors. *Vision Res., 29*(11), 1471-1483.

Smodlaka, H., Khamas, W. A., Palmer, L., Lui, B., Borovac, J. A., Cohn, B. A., & Schmitz, L. (2016). Eye histology and ganglion cell topography of northern elephant seals (*Mirounga angustirostris*). *The Anatomical Record, 299*(6), 798-805. doi:<https://doi.org/10.1002/ar.23342>

Stone, J., & Johnston, E. (1981). The topography of primate retina: A study of the human, bushbaby, and new- and old-world monkeys. *J. Comp. Neurol., 196*(2), 205-223. doi:<https://doi.org/10.1002/cne.901960204>

Tetreault, N., Hakeem, A., & Allman, J. M. (2004). The distribution and size of retinal ganglion cells in *Microcebus murinus*, *Cheirogaleus medius*, and *Tarsius syrichta*: Implications for the evolution of sensory systems in primates. In C. F. Ross & R. F. Kay (Eds.), *Anthropoid Origins: New Visions* (pp. 463-475). Boston, MA: Springer US.

Tiao, Y.-C., & Blakemore, C. (1976). Regional specialization in the golden hamster's retina. *J. Comp. Neurol., 168*(4), 439-457. doi:<https://doi.org/10.1002/cne.901680402>

Vega-Zuniga, T., Medina, F. S., Fredes, F., Zuniga, C., Severín, D., Palacios, A. G., Karten, H. J., & Mpodozis, J. (2013). Does nocturnality drive binocular vision? Octodontine rodents as a case study. *PLoS One, 8*(12), e84199. doi:10.1371/journal.pone.0084199

Veilleux, C. C., & Christopher Kirk, E. (2009). Visual acuity in the cathemeral strepsirrhine *Eulemur macaco flavifrons*. *Am. J. Primatol., 71*(4), 343-352. doi:<https://doi.org/10.1002/ajp.20665>

Wang, H.-H., Gallagher, S. K., Byers, S. R., Madl, J. E., & Gionfriddo, J. R. (2015). Retinal ganglion cell distribution and visual acuity in alpacas (*Vicugna pacos*). *Vet. Ophthalmol., 18*(1), 35-42. doi:<https://doi.org/10.1111/vop.12131>

Welsch, U., Ramdohr, S., Riedelsheimer, B., Hebel, R., Eisert, R., & Plötz, J. (2001). Microscopic anatomy of the eye of the deep-diving Antarctic Weddell seal (*Leptonychotes weddellii*). *J. Morphol., 248*(2), 165-174. doi:<https://doi.org/10.1002/jmor.1027>

Wilder, H. D., Grünert, U., Lee, B. B., & Martin, P. R. (1996). Topography of ganglion cells and photoreceptors in the retina of a New World monkey: The marmoset *Callithrix jacchus*. *Vis. Neurosci., 13*(2), 335-352. doi:10.1017/S0952523800007586

Xiao, X., Zhao, T., Miyagishima, K. J., Chen, S., Li, W., & Nadal-Nicolás, F. M. (2021). Establishing the ground squirrel as a superb model for retinal ganglion cell disorders and optic neuropathies. *Lab. Invest., 101*(9), 1289-1303. doi:<https://doi.org/10.1038/s41374-021-00637-y>
